# Functionally distinct BMP1 isoforms show an opposite pattern of abundance in plasma from non-small cell lung cancer subjects and controls

**DOI:** 10.1101/2022.01.07.475393

**Authors:** Margaret K. R. Donovan, Yingxiang Huang, John E. Blume, Jian Wang, Daniel Hornburg, Shadi Ferdosi, Iman Mohtashemi, Sangtae Kim, Marwin Ko, Ryan W. Benz, Theodore L. Platt, Serafim Batzoglou, Luis A. Diaz, Omid C. Farokhzad, Asim Siddiqui

## Abstract

Advancements in deep plasma proteomics are enabling high-resolution measurement of plasma proteoforms, which may reveal a rich source of novel biomarkers previously concealed by aggregated protein methods. Here, we analyze 188 plasma proteomes from non-small cell lung cancer subjects (NSCLC) and controls to identify NSCLC-associated protein isoforms by examining differentially abundant peptides as a proxy for isoform-specific exon usage. We find four proteins comprised of peptides with opposite patterns of abundance between cancer and control subjects. One of these proteins, BMP1, has known isoforms that can explain this differential pattern, for which the abundance of the NSCLC-associated isoform increases with stage of NSCLC progression. The presence of cancer and control-associated isoforms suggests differential regulation of BMP1 isoforms. The identified BMP1 isoforms have known functional differences, which may reveal insights into mechanisms impacting NSCLC disease progression.

## Introduction

Multiple isoforms of a single protein, or proteoforms, can arise due to alternative splicing (i.e., protein isoforms), allelic variation, and post translational modifications ^1^. Proteoforms play key and distinct roles in biological mechanisms, including impacting complex traits^2^ and disease^3^. For example, protein isoforms may differ in domain composition, where consequently each isoform may have substantially different functions and influence disease predisposition or progression. Advances in characterizing the proteomic landscape of lung cancers such as non-small cell lung cancer (NSCLC) and squamous cell lung cancer have enabled identification of important protein biomarkers^4–6^, however, few proteoforms relevant to lung cancer have been identified^7^, as these studies are limited to only single or few protein^8–10^ or proteoforms arising from different genes^11^. Unbiased readout technologies, such as high-resolution quantitative mass spectrometry (MS), can be employed to infer and quantify peptides and proteins with high confidence (e.g., < 1% false discovery rate (FDR)). However, large-scale LC-MS/MS-based proteomics studies have historically been impractical due to cumbersome and lengthy workflows required to achieve unbiased, deep, and rapid sampling of clinically relevant biospecimens with large dynamic ranges of protein abundances, such as blood plasma ^12–14^.

Here, we analyze data from a previous study^15^ of independent acquisition (DIA)-based MS data generated from 188 subjects (80 healthy control subjects and 108 subjects identified as having NSCLC) using the Proteograph™ workflow which uses nanoparticles (NPs) to enable high-resolution, unbiased, and deep assessment of the plasma proteome. We used a discordant peptide intensity search (Figure 1A) to infer four proteins with differentially abundant protein isoforms, including BMP1, for which we show has differential abundance of two isoforms (long and short), both of which have higher magnitude of differential abundance at later stages of NSCLC. BMP1 plays a role in collagen processing and the short isoform lacks the domains enabling its secretion, potentially impacting collagen’s protective role in cancer consistent with the higher abundance of the short isoform in cancer subjects observed in this paper. Hence, BMP1 isoforms may constitute a novel biomarker previously concealed when assessing the aggregated BMP1 protein abundance.

**Fig 1.**
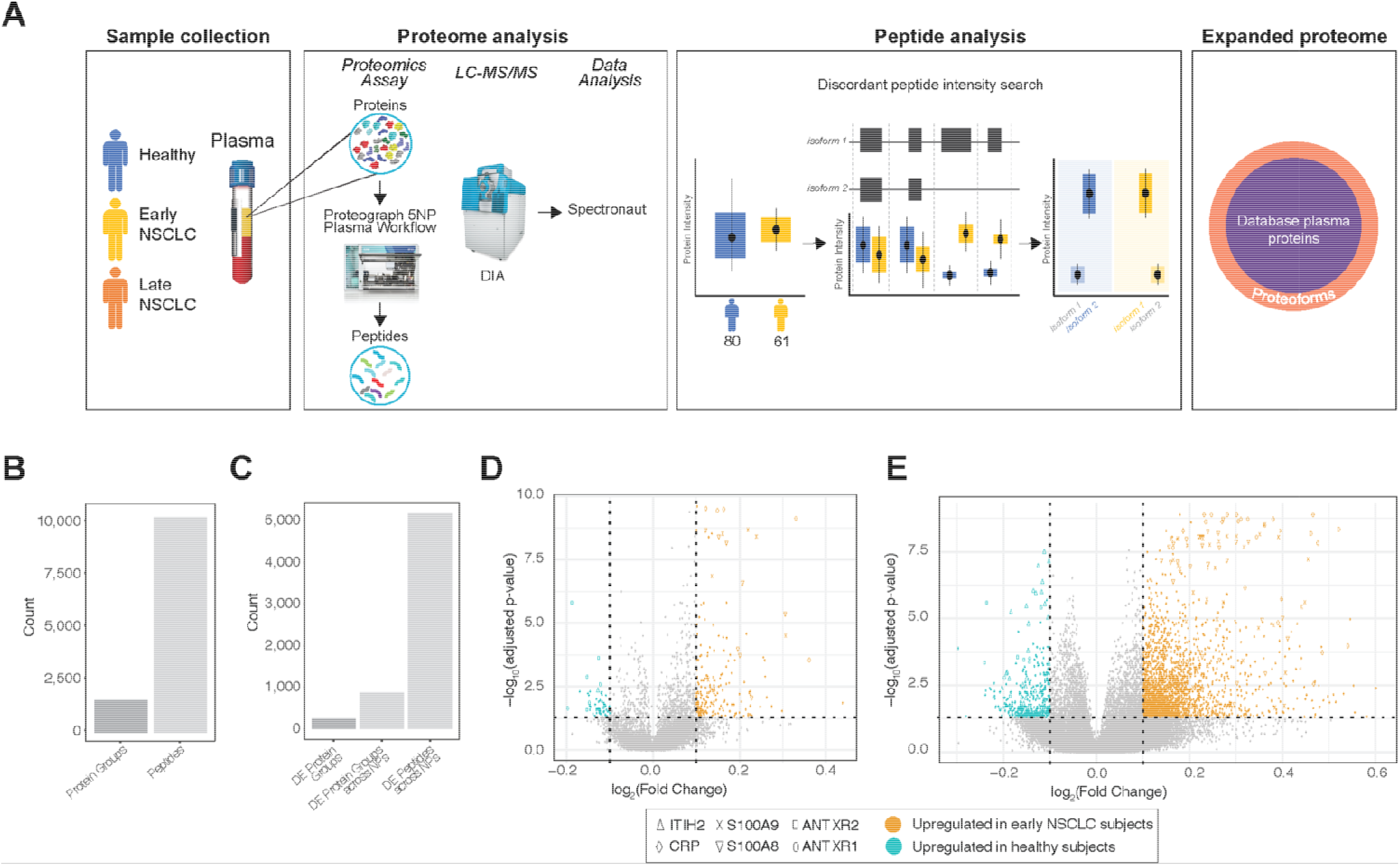
Proteome analysis of healthy and NSCLC subjects using a 5 NP plasma workflow. A. Overview of this proof-of-concept proteoform identification study. Plasma samples were collected from healthy (blue), early non-small cell lung cancer (NSCLC; yellow), late NSCLC (orange), and co-morbid (green) subjects (*Sample Collection*). The plasma proteomes were analyzed for each of these subjects, which included protein extraction, protein discovery using the NP-based Proteograph platform, then DIA protein/peptide identification and quantification using LC-MS/MS and search algorithms (*Proteome Analysis*). Proteoforms were then identified using a discordant peptide intensity search, which included examining peptide mappings to known protein coding isoforms and using differential abundance to discover protein isoforms. Together, these identified proteoforms represent an expanded plasma proteome database not captured in standard MS-based or targeted proteomic studies (*Expanded proteome*). B. Barplots showing the number of peptides and proteins groups retained after filtering to those present in at least 50% of subjects from either heathy or early NSCLC. C. Barplots showing the number of differentially abundant (DA): 1) protein groups, with collapsed abundances using MaxLFQ; 2) protein groups across NPs (i.e., DA independently across NPs); and 3) peptides across NPs. D. Volcano plot showing the significance (adjusted p-value; y-axis) and fold change (x-axis) from calculating the differential abundance of protein groups across NPs between healthy and early NSCLC subjects. Protein groups with a log2(Fold Change) greater or less than 1.0 and adjusted p-value < 0.05 are highlighted, where protein groups with increased abundance in early NSCLC subjects are shown in orange and protein groups with increased abundance in healthy subjects are shown in teal. Proteins with known roles in cancer and immune response (ITIH2, CRP, S100A9, S100A8, ANTXR2, and ANTXR1) are highlighted with various shapes. E. Volcano plot showing the significance (adjusted p-value; y-axis) and fold change (x-axis) from calculating the differential abundance of peptides across NPs between healthy and early NSCLC subjects. Peptides with a log2(Fold Change) greater or less than 1.0 and adjusted p-value < 0.05 are highlighted, where peptides with increased abundance in early NSCLC subjects are shown in orange and peptides with increased abundance in healthy subjects are shown in teal. Peptides mapping to proteins with known roles in cancer and immune response (ITIH2, CRP, S100A9, S100A8, ANTXR2, and ANTXR1) are highlighted with various shapes.

## Results

### Peptide-level analyses provides unique biological insight versus protein-level

Starting from the previously derived analysis^15^, we searched for proteins and peptides that are differentially abundant (DA). First, to reduce potential noise introduced by rare peptides, proteins were filtered to those present in at least 50% of subjects from either 80 healthy or 61 early NSCLC (stages 1, 2 and 3) subjects, retaining 10,280 peptides and 1,565 proteins across 141 subjects (Figure 1B). Next, as each protein may have been detected by more than one NP (each NP can be thought of as generating a separate MS fraction), we use MaxLFQ^16^ to quantify a single abundance (hereto referred to as *collapsed abundances*) between healthy and early NSCLC subjects. We evaluated differential protein abundance observing 243 significantly regulated proteins (adjusted p < 0.05; Wilcoxon Test) (Figure 1C, D). To investigate NPs capacity to capture biological signal beyond abundance levels (e.g., proteoform information, or NP specific protein complexes), we treated each NP:protein feature pair as a separate observation comparing healthy and early NSCLC subjects. We identified 877 NP:protein feature pairs (Figure 1E), corresponding to a 3.6-fold increase from examining differences at the aggregated level alone. This highlights the capacity of NPs coupled with LC-MS/MS to interrogate the proteome at a finer biological resolution (i.e. protein variants and complexes) than that captured by conventional DA analysis at the aggregated protein level. In addition, we performed DA analysis using peptide abundances across all NPs (i.e., not collapsed abundances) between healthy and early NSCLC subjects and identified 5,181 DA peptides (Figure 1C, E), corresponding to a 6.5-fold increase from examining differences at the protein-level. Further, we identified known hallmark cancer and inflammatory biomarkers which were differentially regulated in the peptide data (Supplemental Results, Figure 1D,E). Overall, this increased number of observed significant differences between proteins, protein across NPs, and peptides across NPs, verified by the presence of known cancer biomarkers, indicates substantial opportunity to increase biological insight and the potential to identify proteoforms using peptide-level, high resolution proteomics.

### Identification of four NSCLC-associated proteoforms using peptide-level discordant peptide search

Next, we explored whether we could use DA peptides in contrast to the average protein-level information to help resolve proteoforms. Specifically, we extracted DA peptides and retained proteins with at least one peptide over-expressed in healthy subjects and at least one peptide over-expressed in early NSCLC subjects (Figure 2A). Then, by mapping the DA peptides to genomic space, we inferred potential exon usage and proteoforms. We performed this discordant peptide intensity analysis and identified four proteins for which we potentially captured multiple protein isoforms with significant differential behavior in early NSCLC when compared to healthy controls: BMP1, C4A, C1R, and LDHB (Figure 2B). We examined the Open Target Score ^17^ (Release 21.09), which is an association score of known and potential drug targets with diseases using integrated genome-wide data from a broad range of data sources, to assess the association of the four proteins with lung carcinoma targets. We found modest to low scores (Figure 2B), suggesting a mix of novel and known lung cancer-associated proteins. These proteins have all been previously identified in plasma and range from highly abundant (C4A, C1R, LDHB) to moderately abundant (BMP1) ^18^ (Figure 2C). BMP1, the least abundant of the four proteins, is not identified in depleted plasma published in this study., indicating this approach identified protein isoforms inaccessible with conventional depleted plasma proteomics workflows. These results indicate that, using a MS-based peptide discordant intensity search, we can infer proteoforms with possible relevance to NSCLC.

**Fig 2.**
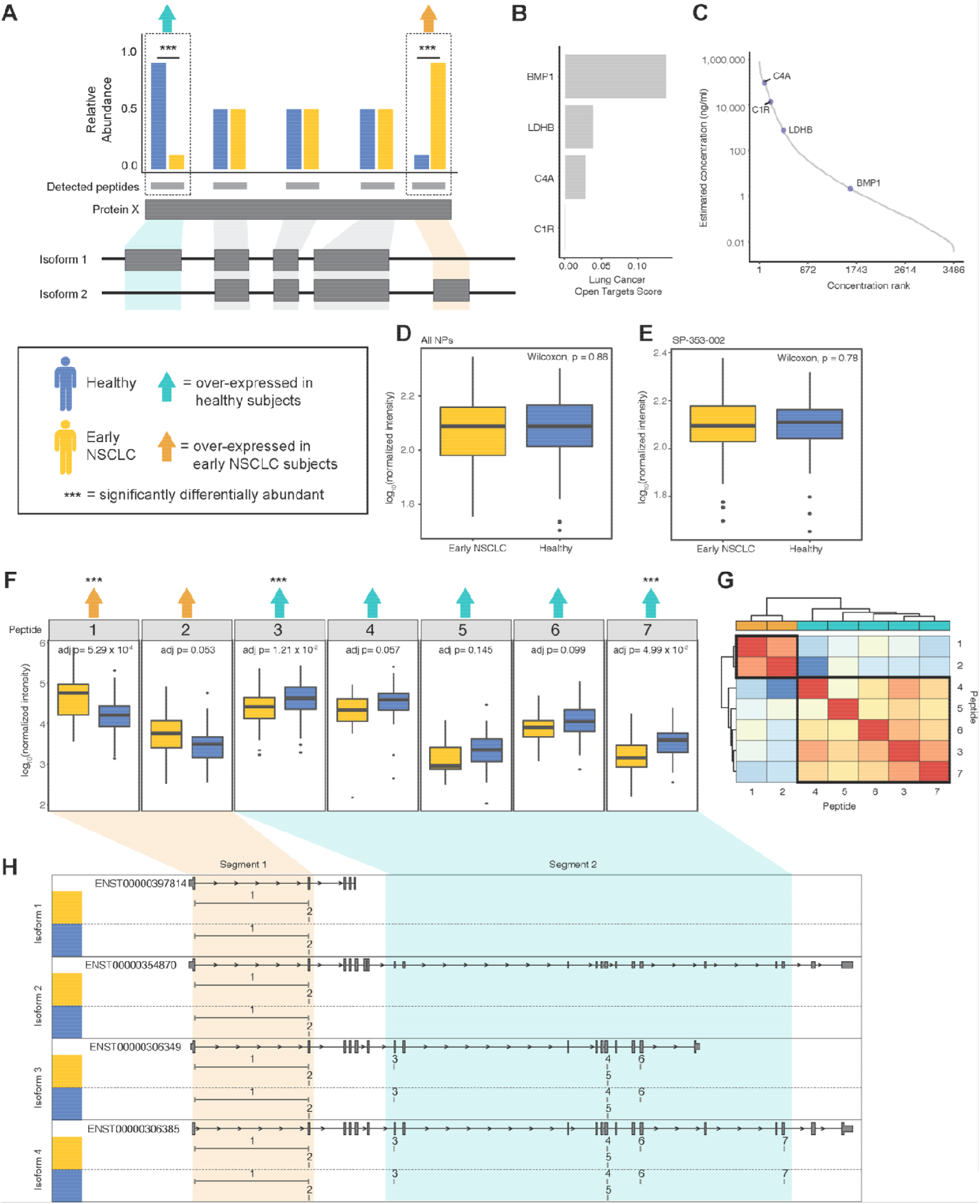
Identification of four proteoforms, including BMP1, in 141 healthy and early NSCLC subjects using a discordant peptide intensity search. A. Cartoon describing the discordant peptide intensity search strategy. We calculated DA across peptides between healthy (blue) and early NSCLC (yellow). Protein groups with at least one peptide significantly over-expressed (triple asterisks) in healthy subjects (teal arrow) and at least one peptide over-expressed in early NSCLC subjects (orange arrow) were identified as having putative proteoforms. Mapping the peptides to the gene structure, we inferred potential exon usage and segments suggesting the detection of more than one protein isoform. B. Barplot showing four proteins in which we potentially captured multiple protein isoforms: BMP1, C4A, C1R, and LDHB and their associated Open Target Score for lung carcinoma. C. Plot showing the four proteins with putative proteoforms matched to a reference database (HPPP) plotted as a distribution by the rank order of published concentrations (x-axis) and by the log_10_ published concentration (ng/ml; y-axis). D. Box plot showing the log_10_ median normalized intensities of BMP1 in early NSCLC subjects (yellow) and in healthy subjects (blue) with collapsed abundances across NPs. P-values, calculated using a Wilcoxon test, are shown. E. Box plot showing the log_10_ median normalized intensities of BMP1 in early NSCLC subjects (yellow) and in healthy subjects (blue) in NP, SP-353-002. P-values, calculated using a Wilcoxon test, are shown. F. Series of boxplots showing the log_10_ median normalized intensities of seven peptides mapping BMP1 in early NSCLC (yellow) and healthy subjects (blue). Peptides that are over-expressed in healthy subjects are indicated with a teal arrow and in early NSCLC are indicated with an orange arrow. Peptides that are significantly DA are indicated with a triple asterisk. P-values, calculated using a Wilcoxon test and adjusted, are shown. G. Heatmap showing the Pearson correlation of the seven BMP1 peptide abundances, where low correlation is indicated in shades of blue and high correlation is indicated in shades of red. Correlation values were clustered using hierarchical clustering. Peptides are annotated by the direction of DA, including over-expressed in healthy subjects are highlighted in teal and early NSCLC are highlighted in orange. H. Gene structure plots of four known BMP1 protein coding transcripts (i.e., isoforms) with the seven BMP1 peptides mapped to genomic region. Peptides spanning intronic regions are indicated with a horizontal line. Peptides 1 and 2, corresponding to being over-expressed early NSCLC, are boxed in orange, creating one segment. Peptides 37, corresponding to being over-expressed healthy, are boxed in teal, creating a second segment. Segment 2 appears to correspond to the shorter isoform 1, whereas segment 2 appears to correspond to the longer isoforms 2-4.

To interrogate the extent to which isoforms information adds to disease insight, we examined differences in abundances between healthy and early NSCLC subjects for BMP1 (Figure 2D-G), C4A (Supplemental Figure 1), C1R (Supplemental Figure 2), and LDHB (Supplemental Figure 3) at the collapsed protein-level, NP:protein-level, and peptide-level. Examining BMP1, at the collapsed protein (Figure 2D) and NP:protein (Figure 2E) level, we do not observe a difference in BMP1 abundance, as a result of an averaging of peptide abundances occurring at the protein-level. However, at the peptide-level (Figure 2F), there are three significantly differential peptides: 1) peptide 1, which is significantly upregulated in early NSCLC subjects (adjusted p = 5.29 × 10^−4^; Wilcoxon Test); 2) peptide 3, which is significantly upregulated in healthy subjects (adjusted p = 1.21 × 10^−2^; Wilcoxon Test); and 3) peptide 7, which is significantly upregulated in healthy subjects (adjusted p = 4.99 × 10^−2^; Wilcoxon Test). We also observe a trend in direction of abundance differences, where the first two peptides are upregulated in early NSCLC subjects and the last five peptides are upregulated in healthy subjects (Figure 2F). To assess whether these two groups of peptides belong to different proteoforms, we further compared their abundance similarities across the 141 subjects. We expect peptides that belong to the same proteoform to have correlated abundances across a cohort of individuals since they belong to the same molecular entity while peptides belonging to different proteoforms should have less-correlated abundances across the same cohort of individuals. We thus performed pairwise Pearson correlation and hierarchical clustering analysis, which showed two distinct clusters driven by a high degree of correlation in peptide 1 and 2 (cluster 1) and peptides 3-7 (cluster 2) (Figure 2G). We next mapped the peptides to their genomic sequence, including four protein coding isoform transcripts (ENST00000397814, ENST00000354870, ENST00000306349, and ENST00000306385), and ordered them according to exon order (Figure 2H). We observed two distinct segments of corresponding direction of BMP1 peptide differential abundance. Specifically, peptides 1 and 2 were both upregulated in early NSCLC subjects (segment 1) and peptide 3-7 were all upregulated in healthy subjects (segment 2) (Figure 2F). Peptides mapping segment 1 exclusively map to exons present in the short isoform (ENST00000397814), whereas peptides mapping to segment 2 exclusively map to exons present in the three longer isoforms (ENST00000354870, ENST00000306349, and ENST00000306385) (Figure 2H). The opposite pattern of abundance of the long and short isoforms in early NSCLC subjects versus healthy subjects suggest that BMP1 isoforms may play a role in cancer and may serve as a novel biomarker‥ This pattern is exaggerated when examining long and short isoforms in late-stage NSCLC subjects, where we observe a trend of increasing upregulation of segment 1 peptides (short BMP1 isoform) and a trend of decreasing upregulation of segment 2 peptides (long BMP1 isoform) between healthy subjects, Stage 1 and 2 NSCLC subjects, and Stage 3 and 4 NSCLC subjects (Supplemental Figure 4).

## Discussion

Existing technologies, including an unbiased bottom-up NP-based methodology upstream of LC-MS/MS-based workflows and targeted methodologies, have enabled protein-centric analyses that have revealed new insights into human disease. While protein-centric bottom-up analyses have made substantial strides in our understanding of human biology, aggregating peptide level quantifications to the protein level may conceal biologically critical features, such as proteoforms arising from alternative splicing (protein isoform), allelic variation (protein variants), or post-translational modifications, which may provide mechanistic insights and novel biomarkers underlying complex traits and disease. Importantly, unbiased LC-MS/MS-based proteomic data can be re-mined to enable peptide-centric analyses that may reveal new information about proteoforms. In this study, we use peptide-level information derived from LC-MS/MS data to enable proteoform identification using discordant peptide abundance and apply that to a NSCLC cohort. Typically, protein inference engines use peptide-level data to detect the presence or absence of peptides to identify protein isoforms. However, here we show the utility of incorporating quantitative profiles of peptides mapping to known isoforms in potentially increasing the sensitivity of the underlying proteoform detection. Thus, we hypothesized that previously generated LC-MS/MS plasma proteomic data can be reanalyzed at the peptide-level using quantitative profiles to infer protein isoforms^15^, potentially yielding deeper insights into disease mechanisms and we demonstrated that such a reanalysis revealed known and putative, novel disease-relevant proteoforms.

We performed peptide analysis using DIA data derived from healthy and early NSCLC subjects by conducting a discordant peptide intensity search to identify protein isoforms. We identified four proteins with DA peptides and putative isoforms, including BMP1, C4A, C1R, and LDHB. Importantly, none of these proteins showed a difference in abundance at the protein-level. For BMP1 and C1R, using peptide abundance as a proxy for functionally relevant proteins we identified potential NSCLC-related isoforms. We showed BMP1 has differential abundance of two isoforms (long and short), both of which have higher magnitude of differential abundance at later stages of NSCLC. BMP1 plays a role in collagen processing and the short isoform lacks the domains enabling its secretion, potentially impacting collagen’s protective role in cancer consistent with the higher abundance of the short isoform in cancer subjects observed in this paper. Additionally, C4A showed distinct peptide abundance discordance in one segment of the protein, which did not correspond to any known protein coding isoforms, suggesting peptide-centric proteoform identification may result in novel disease-associated isoforms.

The method we used to search for protein isoforms through discordant peptide intensity is stringent in terms of the number of protein isoform candidates we can find, but easily interpretable. Similar approaches such as COPF ^19^ and PeCorA ^20^ use quantitative disagreements between peptides mapped to the same protein or peptide correlation within the same protein to detect protein isoforms and suggest proteoforms. However, as shown with our examples where 2 of the 4 isoform candidates (C1R and LDHB) met the discordant peptide intensity criteria but failed to be readily explained by known isoforms or biological conjecture, evaluation of the validity of the isoform candidate is needed but is outside the scope of this study. In this paper, our validation was mapping the peptides back to the genomic sequence and known isoform transcripts. Manual validation (e.g., isoform specific enrichment with isoform specific antibodies) can confirm the presence of novel isoforms. This might be possible for the limited candidates arising from our stringent isoform detection process, however, other processes such COPF and PeCorA could yield dramatically more candidates.

It is possible that the finding of only four protein isoforms in 188 subjects has been impacted by limited sample sizes reducing our power to identify proteoforms. Similarly, it is also possible that other undiscovered proteoforms are not functional in plasma and may only be identified in other biofluids or tissues. While our study shows the utility of using NP-based methodology upstream LC-MS/MS-based workflows to identify proteoforms, it is possible that expanding the sample size and diversity in sample type may yield further insights into disease-associated proteoforms. In addition, LC-MS/MS enables quantifying and identifying tens of thousands of peptides with post-translational modifications precisely defined by their intact mass and fragmentation pattern.

The identification of proteoforms (protein isoforms) highlights important considerations for current approaches characterizing the impact of genetic variation on molecular phenotypes, like protein abundance, by conducting protein quantitative trait analyses (pQTLs). Specifically, recent pQTL analyses using large cohorts ^21^ are performed at the protein-level and largely miss or misattribute peptide-level proteoform effects. Furthermore, these studies utilize aptamer and antibody-based methodologies that, as been recently shown ^21^, can lead to false discoveries and uncertain identification error rates because of conceptual limitations (e.g., the presence of a non-synonymous SNP inducing an amino acid change that disrupts the binding of the aptamer or antibody). Interrogating protein abundances with this high resolution approach provides deeper insight into the molecular mechanisms underlying human biology and opens a possible new avenue for biomarker identification and therapeutic development.

## Materials and methods

### Identification of protein isoforms

As previously reported, plasma from healthy subjects and from subjects diagnosed with NSCLC at stage 1, 2, 3, and 4 was collected and processed with the Proteograph™ workflow ^15^ and DIA data was generated and processed (Supplemental Methods). From the 1,565 proteins present after filtering, we searched for peptides that had differential abundance between controls and cancer (p < 0.05; Benjamini-Hochberg corrected). Discordant pairs are defined as peptides from the same protein where at least one peptide was identified with significantly higher, and another peptide was identified with significantly lower plasma abundance in healthy controls vs. early NSCLC.

This work generated no additional data from new or existing patient samples and used raw data already deposited in the public database ProteomeXchange Consortium (http://proteomecentral.proteomexchange.org) with the dataset PXD017052.

### Protein quantification across multiple samples

Within each nanoparticle, standard MaxLFQ was used to quantify abundance at the protein level. For each peptide, the intensity ratios between every pair of samples were first computed. The pairwise protein ratio is then defined as the median of the peptide ratios from all peptides map to the same protein. With all the pairwise protein ratios between any two samples, we can perform a least-squares analysis to reconstruct the abundance profile optimally satisfying all the protein ratios. Then the whole profile is rescaled to the cumulative intensity across samples for the final protein abundance ^22^. A modified MaxLFQ was used to quantify abundance across samples and nanoparticles. For each protein, all peptides’ intensities belonging to a protein from all samples and NP were employed to calculate peptide ratios and subsequent calculation steps resulting in abundance across all samples and NP.

## Acknowledgements

We thank the individuals who took part in this study for their participation. We also thank Steven A. Carr and Robert Langer for their review of this manuscript.

## Author contributions

MKRD and YH contributed equally. MKRD, YH, AS, JB, DH, SB, LAD, and OF conceived this study. MKRD and YH prepared the manuscript. AS edited the manuscript. MK, RB, TP processed the DIA data. YH, AS, JB, and MKRD performed the protein isoform analyses. IM, SF, and DH supported LC-MS/MS data analyses. All authors discussed and reviewed the manuscript.

## References

1. Smith, L. M., Kelleher, N. L. & Proteomics, T. C. for T. D. Proteoform: a single term describing protein complexity. Nat Methods 10, 186–187 (2014).

2. Li, Y. I. et al. RNA splicing is a primary link between genetic variation and disease. Science (80-.). 352, 600–604 (2016).

3. Lisitsa, A., Moshkovskii, S., Chernobrovkin, A., Ponomarenko, E. & Archakov, A. Profiling proteoforms: promising follow-up of proteomics for biomarker discovery. Expert Rev. Proteomics 11, 121–129 (2014).

4. Satpathy, S. et al. A proteogenomic portrait of lung squamous cell carcinoma. Cell 184, 4348–4371.e40 (2021).

5. Kisluk, J., Ciborowski, M., Niemira, M., Kretowski, A. & Niklinski, J. Proteomics biomarkers for non-small cell lung cancer. J. Pharm. Biomed. Anal. 101, 40–49 (2014).

6. Nishimura, T., Végvári, Á., Nakamura, H., Kato, H. & Saji, H. Mutant Proteomics of Lung Adenocarcinomas Harboring Different EGFR Mutations. Front. Oncol. 10, 1–20 (2020).

7. Nishimura, T. et al. Current status of clinical proteogenomics in lung cancer. Expert Rev. Proteomics 16, 761–772 (2019).

8. Franc, V., Zhu, J. & Heck, A. J. R. Comprehensive Proteoform Characterization of Plasma Complement Component C8αβγ by Hybrid Mass Spectrometry Approaches. J. Am. Soc. Mass Spectrom. 29, 1099 (2018).

9. Gao, J. et al. Within-person reproducibility of proteoforms related to inflammation and renal dysfunction. Sci. Rep. 12, (2022).

10. Koska, J. et al. Plasma proteoforms of apolipoproteins C-I and C-II are associated with plasma lipids in the Multi-Ethnic Study of Atherosclerosis. J. Lipid Res. 63, 100263 (2022).

11. Wåhlén, K. et al. Significant correlation between plasma proteome profile and pain intensity, sensitivity, and psychological distress in women with fibromyalgia. Sci. Rep. 10, 12508 (2020).

12. Anderson, N. L. The clinical plasma proteome: a survey of clinical assays for proteins in plasma and serum. Clin. Chem. 56, 177–185 (2010).

13. Geyer, P. E., Holdt, L. M., Teupser, D. & Mann, M. Revisiting biomarker discovery by plasma proteomics. Mol. Syst. Biol. 13, 942 (2017).

14. Geyer, P. E. et al. Plasma Proteome Profiling to Assess Human Health and Disease. Cell Syst. 2, 185–195 (2016).

15. Blume, J. E. et al. Rapid, deep and precise profiling of the plasma proteome with multi-nanoparticle protein corona. Nat. Commun. 11, 1–14 (2020).

16. Zhu, Y. et al. DEqMS: A method for accurate variance estimation in differential protein expression analysis. Mol. Cell. Proteomics 19, 1047–1057 (2020).

17. Koscielny, G. et al. Open Targets: A platform for therapeutic target identification and Validation. Nucleic Acids Res. 45, D985–D994 (2017).

18. Schwenk, J. M. et al. The Human Plasma Proteome Draft of 2017: Building on the Human Plasma PeptideAtlas from Mass Spectrometry and Complementary Assays. J Proteom Res 16, 4299–4310 (2017).

19. Bludau, I. et al. Systematic detection of functional proteoform groups from bottom-up proteomic datasets. Nat. Commun. 12, (2021).

20. Dermit, M., Peters-Clarke, T. M., Shishkova, E. & Meyer, J. G. Peptide Correlation Analysis (PeCorA) Reveals Differential Proteoform Regulation. J. Proteome Res. 20, 1972–1980 (2021).

21. Pietzner, M. et al. Synergistic insights into human health from aptamer- and antibody-based proteomic profiling. Nat. Commun. 12, (2021).

22. Cox, J. et al. Accurate proteome-wide label-free quantification by delayed normalization and maximal peptide ratio extraction, termed MaxLFQ. Mol. Cell. Proteomics 13, 2513–2526 (2014).

