## Supplemental_Material for "Functionally distinct BMP1 isoforms show an opposite pattern of abundance in plasma from non-small cell lung cancer subjects and controls"

### Supporting Information

#### SUPPLEMENTARY RESULTS

#### High-resolution proteomics identifies known and novel biomarkers

To test the utility of NP and peptide-level interrogation of complex biological samples, we examined the top 10 most DA proteins and peptides. Among the top DA peptides, we observed peptides mapping to ITIH2, ANTXR2, and ANTXR1, which are known to be downregulated in early NSCLC plasma samples. Downregulation of ITIH2 expression has been seen in 70% of breast cancers, 71% of lung cancers, and 70% of renal tumors ^1^. ANTXR2/CMG2 was shown to inhibit breast cancer cell growth and is inversely correlated with disease progression and prognosis ^2^. ANTXR1 can reduce tumor growth in vivo by targeting cancer stem cells in conjunction with LeTx ^3^. In agreement with results of other studies, our analysis of NSCLC plasma samples showed upregulation of well-defined pro-inflammatory and cancer biomarkers such as CRP, S100A9, and S100A8 ^4,5^. Together, the observation of known hallmark cancer and inflammatory biomarkers indicates Proteograph-derived proteomic data captures known biological differences and may suggest the presence of other novel biomarkers.

##### In-depth examination of peptides from C4A, C1R, and LDHB isoforms

To interrogate the potential different isoforms, we next examined C4A. At the collapsed protein (Supplemental Figure 1A) and NP:protein (Supplemental Figure 1B) level, as for BMP1, the difference in C4A abundance is not statistically significant, however at the peptide level there are many significantly differentially expressed peptides (Supplemental Figure 1D). Like BMP1, this further supports protein abundance comparisons mask differences that occur at the peptide level. Pairwise Pearson correlation and hierarchical clustering analysis showed two distinct clusters driven by peptide abundance correlation in peptides 40-54 (cluster 1) and peptides 1-39 and 55-64 (cluster 2) (Supplemental Figure 1C). Mapping the peptides to the two known protein coding isoform transcripts (ENST00000428956 and ENST00000498271) and ordering them according to exon order (Supplemental Figure 1E), we observed three distinct segments of corresponding direction of C4A peptide differential expression abundance. Specifically, peptides 1-39 were upregulated in healthy subjects (segment 1), peptides 40-54 were upregulated in early NSCLC subjects (segment 2), and peptides 55-64 were upregulated in healthy subjects (segment 3) (Supplemental Figure 1D) (Table S6). Interestingly, segments 1 and 3 correspond to cluster 2 and segment 2 corresponds to cluster 1, indicating discordant expression of NSCLC-associated exons in segment 2. In contrast to results from analysis BMP1, there was no obvious association with the two known C4A protein coding isoform transcripts. However, the discordant peptides of segment 2 suggest the presence of an unknown isoform or a smaller byproduct from C4A or other members of the C4 complex. ^6^

To interrogate the other potential isoforms, we next examined C1R and LDHB. At the collapsed protein (Supplemental Figure 2A, 3A) and NP:protein (Supplemental Figure 2B, 3B) level, the differences in C1R and LDHB abundances, respectively, were not statistically significant. However, at the peptide level we observe differentially expressed peptides (Supplemental Figure 2D, 3D), indicating protein abundance comparisons mask differences that occur at the peptide level. Pairwise Pearson correlation and hierarchical clustering analysis showed one distinct cluster in each protein. For C1R, this consisted of moderately correlated peptides 1, 2, 4, 6-11, 14, 16, and 17 (cluster 1) and a weak correlation between peptides 3, 5, 12, 13, and 15 (Supplemental Figure 2C). For LDHB, this consisted of highly correlated peptides 1-3, 6, and 8-11 (cluster 1) and a weak correlation between peptides 4, 5, and 7 (Supplemental Figure 3C).

We mapped the peptides to the known protein coding isoform transcripts (C1R: ENST00000647956, ENST00000536053, ENST00000535233, ENST00000649804, ENST00000543835, and ENST00000540242; and LDHB: ENST00000647956, ENST00000536053, ENST00000535233, ENST00000649804, ENST00000543835, and ENST00000540242) and ordered them according to exon order (Supplemental Figure 2E, 3E). There was no clear pattern in healthy or NSCLC subject peptide upregulation corresponding to any of the known isoforms for either protein. However, we did observe upregulation in C1R peptides 14-17 in healthy subjects corresponding to the two short isoforms (isoforms 5 and 6; Supplemental Figure 2E). Beyond this observation, we could not explain the discordance in peptide abundances. Further examination of C1R and LDHB protein isoforms is needed to further explain the discordance in peptide abundances. Together, these results indicate discordant peptide abundance can be utilized to identify some disease-relevant protein isoforms, as was observed in the case of BMP1 and C4A. However, this approach cannot explain all discordance, as was observed in the case of C1R and LDHB. Further work may expand our understanding of disease-associated proteoforms.

#### SUPPLEMENTARY FIGURES

#### Figure S1: Identification of C4A proteoform in in 141 healthy and early NSCLC subjects using a discordant peptide intensity search

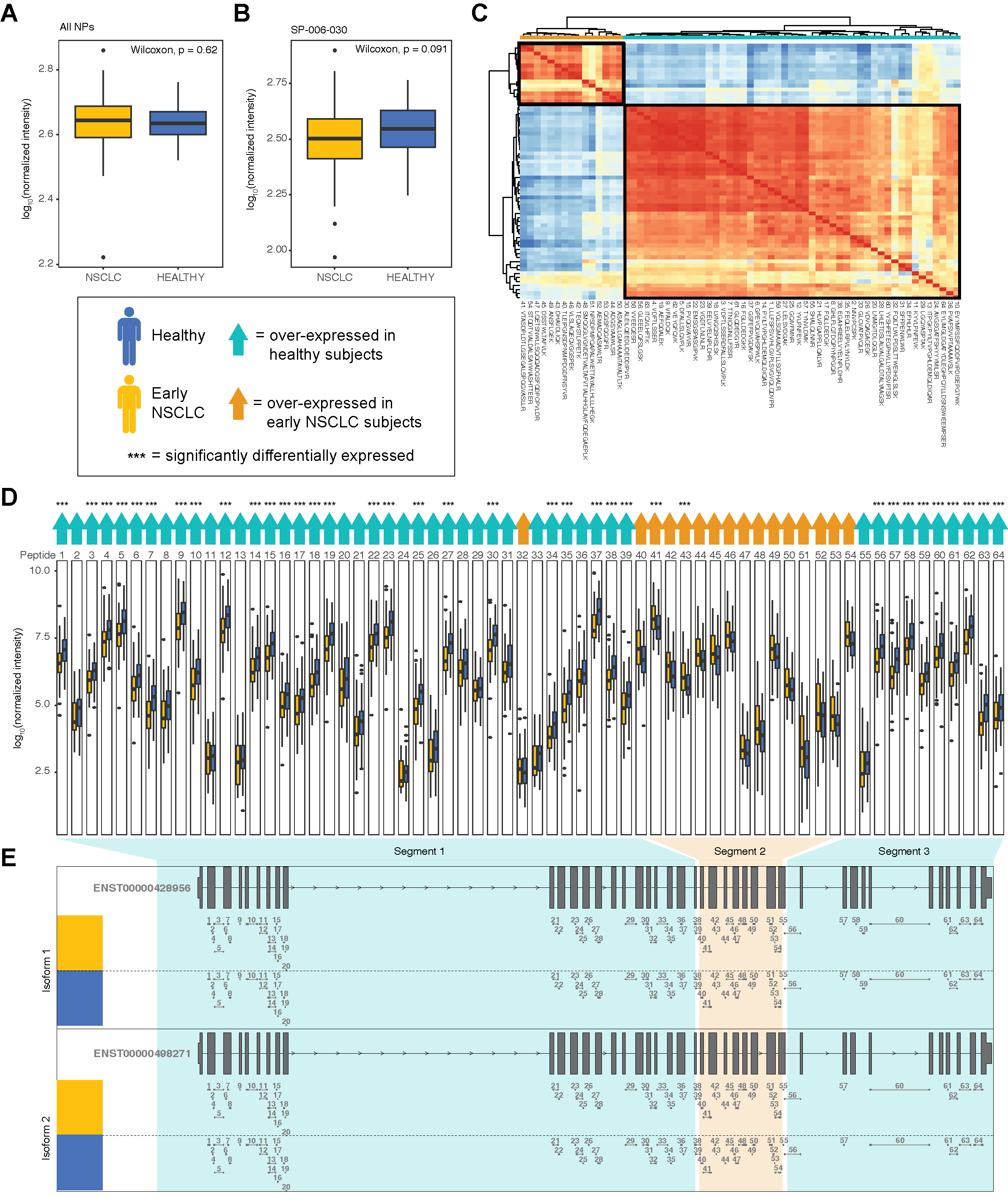

A. Box plot showing the log_10_ median normalized intensities of C4A in early NSCLC subjects (yellow) and in healthy subjects (blue) with collapsed abundances across NPs. P-values, calculated using a Wilcoxon test, are shown.
B. Box plot showing the log_10_ median normalized intensities of C4A in early NSCLC subjects (yellow) and in healthy subjects (blue) in NP, SP-006-030. P-values, calculated using a Wilcoxon test, are shown.
C. Heatmap showing the Pearson correlation of the 64 C4A peptide abundances, where low correlation is indicated in shades of blue and high correlation is indicated in shades of red. Correlation values were clustered using hierarchical clustering. Peptides are annotated by the direction of DE, including over-expressed in healthy subjects are highlighted in teal and early NSCLC are highlighted in orange.
D. Series of boxplots showing the log_10_ median normalized intensities of 64 peptides mapping C4A in early NSCLC (yellow) and healthy subjects (blue). Peptides that are over-expressed in healthy subjects are indicated with a teal arrow and in early NSCLC are indicated with an orange arrow. Peptides that are significantly DE are indicated with a triple asterisk. P-values, calculated using a Wilcoxon test and adjusted, are shown.
E. Gene structure plots of 2 known C4A protein coding transcripts (i.e., isoforms) with the 64 C4A peptides mapped to genomic region. Peptides spanning intronic regions are indicated with a horizontal line. Peptides 1-39 (except peptide 32), corresponding to being over-expressed in healthy subjects, are boxed in teal, creating one segment. Peptides 40-53, corresponding to being over-expressed early NSCLC, are boxed in orange, creating a second segment. Peptides 54-63 (except peptide 64), corresponding to being over-expressed in healthy subjects, are boxed in teal, creating a third segment. Segment patterns do not appear to correspond to any known protein isoforms, potentially indicating a novel isoform.

#### Figure S2: Identification of C1R proteoform in in 141 healthy and early NSCLC subjects using a discordant peptide intensity search

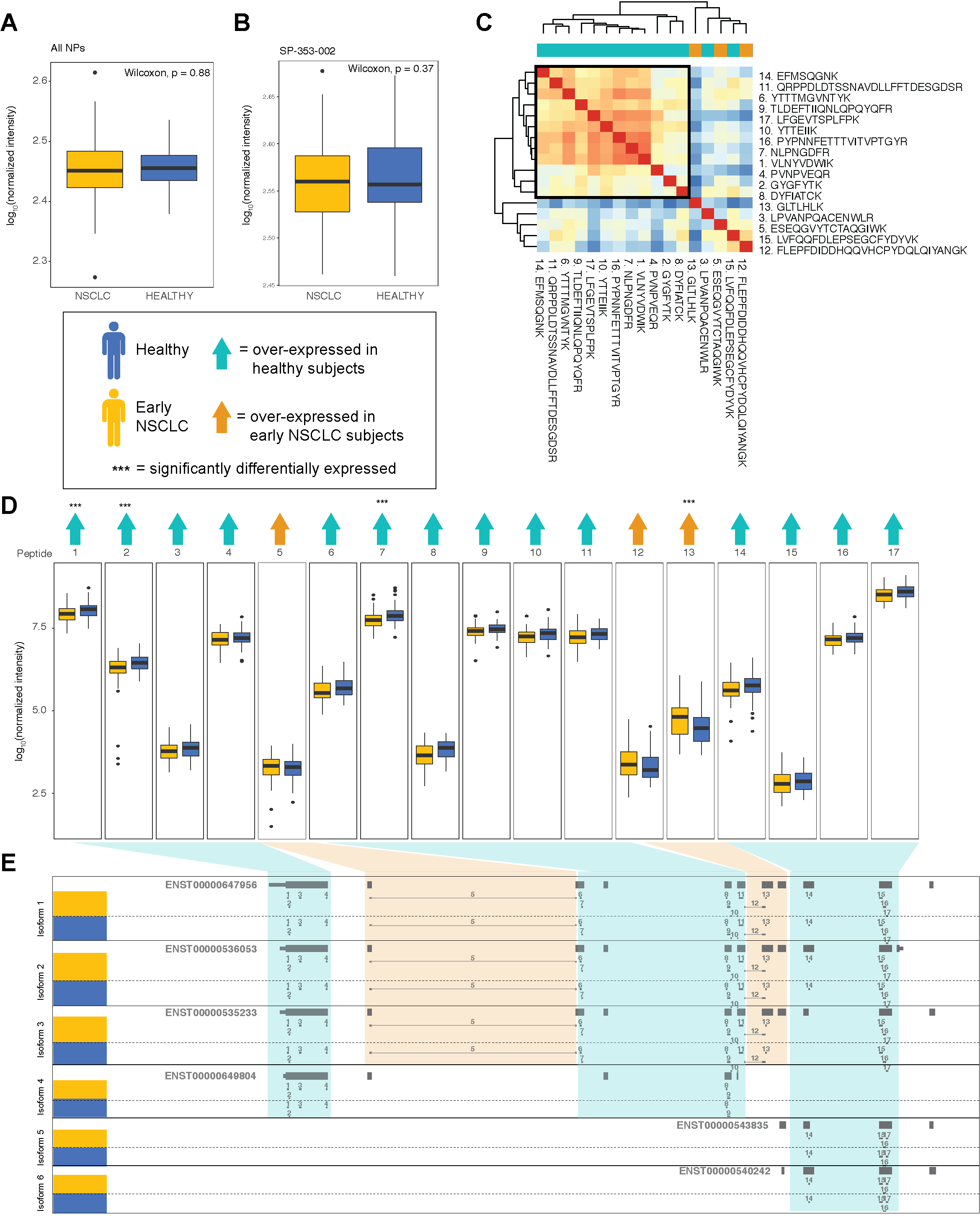

A. Box plot showing the log_10_ median normalized intensities of C1R in early NSCLC subjects (yellow) and in healthy subjects (blue) with collapsed abundances across NPs. P-values, calculated using a Wilcoxon test, are shown.
B. Box plot showing the log_10_ median normalized intensities of C1R in early NSCLC subjects (yellow) and in healthy subjects (blue) in NP, SP-353-002. P-values, calculated using a Wilcoxon test, are shown.
C. Heatmap showing the Pearson correlation of the 17 C1R peptide abundances, where low correlation is indicated in shades of blue and high correlation is indicated in shades of red. Correlation values were clustered using hierarchical clustering. Peptides are annotated by the direction of DE, including over-expressed in healthy subjects are highlighted in teal and early NSCLC are highlighted in orange.
D. Series of boxplots showing the log_10_ median normalized intensities of 17 peptides mapping C1R in early NSCLC (yellow) and healthy subjects (blue). Peptides that are over-expressed in healthy subjects are indicated with a teal arrow and in early NSCLC are indicated with an orange arrow. Peptides that are significantly DE are indicated with a triple asterisk. P-values, calculated using a Wilcoxon test and adjusted, are shown.
E. Gene structure plots of 6 known C1R protein coding transcripts (i.e., isoforms) with the 17 C1R peptides mapped to genomic region. Peptides spanning intronic regions are indicated with a horizontal line. Peptides corresponding to being over-expressed in healthy subjects are boxed in teal. Peptides corresponding to being over-expressed early NSCLC are boxed in orange.

#### Figure S3: Identification of LDHB proteoform in in 141 healthy and early NSCLC subjects using a discordant peptide intensity search

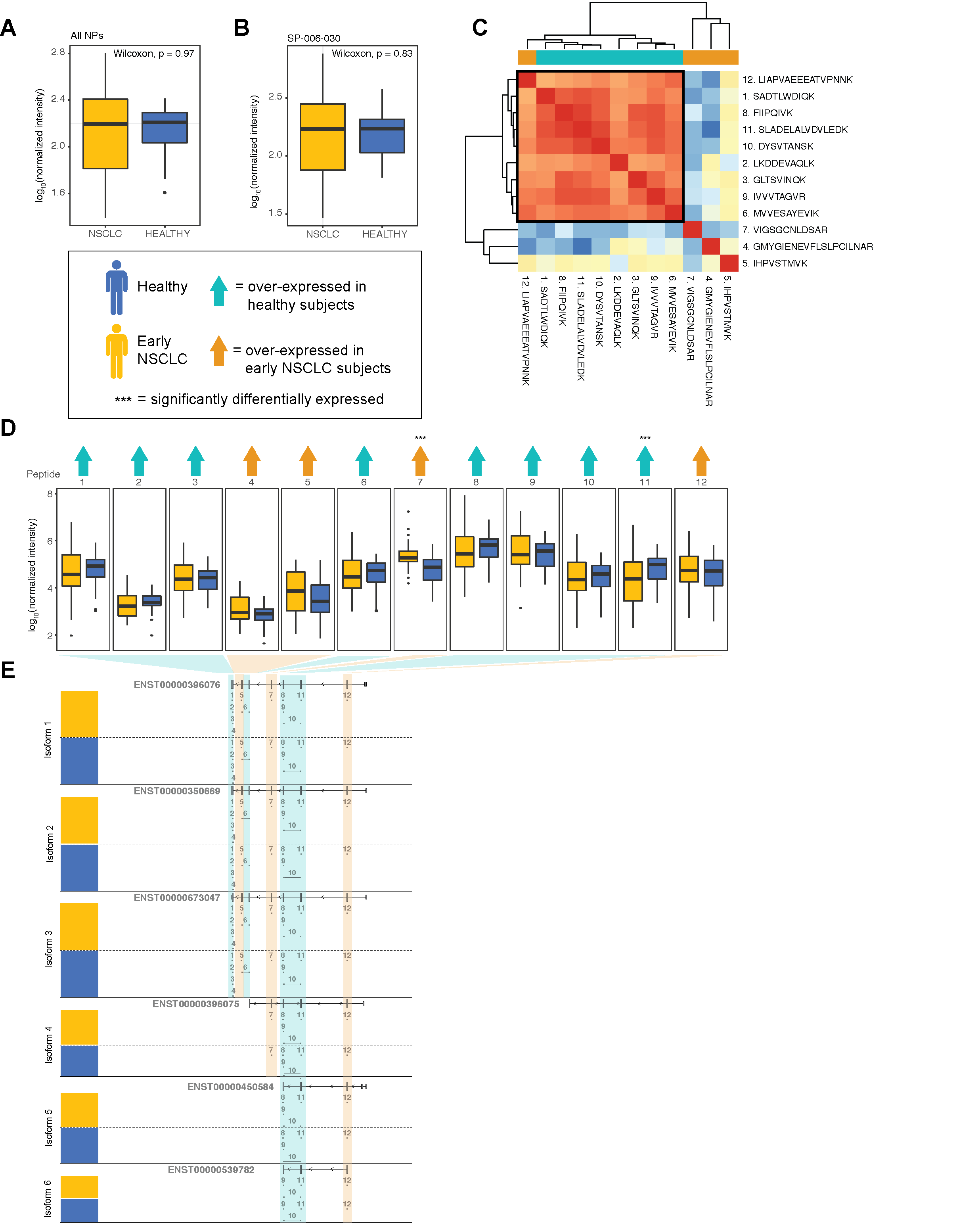

A. Box plot showing the log_10_ median normalized intensities of LDHB in early NSCLC subjects (yellow) and in healthy subjects (blue) with collapsed abundances across NPs. P-values, calculated using a Wilcoxon test, are shown.
B. Box plot showing the log_10_ median normalized intensities of LDHB in early NSCLC subjects (yellow) and in healthy subjects (blue) in NP, SP-006-030. P-values, calculated using a Wilcoxon test, are shown.
C. Heatmap showing the Pearson correlation of the 12 LDHB peptide abundances, where low correlation is indicated in shades of blue and high correlation is indicated in shades of red. Correlation values were clustered using hierarchical clustering. Peptides are annotated by the direction of DE, including over-expressed in healthy subjects are highlighted in teal and early NSCLC are highlighted in orange.
D. Series of boxplots showing the log_10_ median normalized intensities of 12 peptides mapping LDHB in early NSCLC (yellow) and healthy subjects (blue). Peptides that are over-expressed in healthy subjects are indicated with a teal arrow and in early NSCLC are indicated with an orange arrow. Peptides that are significantly DE are indicated with a triple asterisk. P-values, calculated using a Wilcoxon test and adjusted, are shown.
E. Gene structure plots of 6 known LDHB protein coding transcripts (i.e., isoforms) with the 12 LDHB peptides mapped to genomic region. Peptides spanning intronic regions are indicated with a horizontal line. Peptides corresponding to being over-expressed in healthy subjects are boxed in teal. Peptides corresponding to being over-expressed early NSCLC are boxed in orange.

**Figure S4. Association of NSCLC stage (1, 2, 3, and 4) and the presence of BMP1 proteoforms using a discordant peptide intensity search**

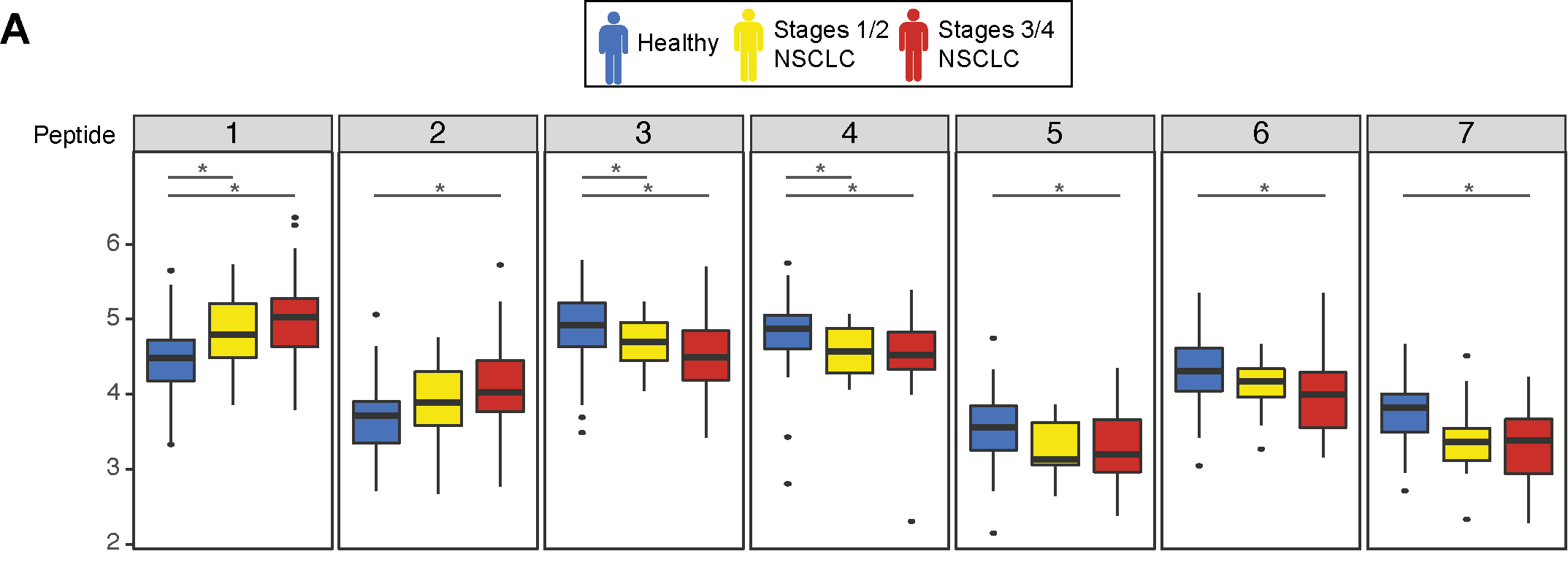

Series of boxplots showing the log_10_ median normalized intensities of 7 peptides mapping BMP1 in healthy subjects (blue, stages 1 and 2 NSCLC (yellow), and stages 3 and 4 (red).

*Peptides that are significantly DE (Wilcoxon test and adjusted).

#### Supplementary Methods

Sample collection and data generation as previously described^7^. Subjects diagnosed with NSCLC stage 1, 2, and 3 were labeled as early NSCLC. Subjects with NSCLC stage 4 were labeled as Late NSCLC. In addition, we have healthy and pulmonary comorbid control arms. Subjects diagnosed with NSCLC but with Unknown stage were removed from analysis; subjects who did not have peptides detected in all nanoparticles in the 10-NP panels were also removed. Summary statistics of protein counts and peptide counts per protein were calculated at this point.

Next, proteins were filtered to those present in at least 50% of subjects from either heathy or early cases, leaving us with a total of 188 subjects (80 control and 108 NSCLC). Peptide intensities were median normalized and natural logged.

#### Supplementary References

1.	Hamm, A. *et al.* Frequent expression loss of Inter-alpha-trypsin inhibitor heavy chain (ITIH) genes in multiple human solid tumors: A systematic expression analysis. *BMC Cancer* **8**, 1–15 (2008).

2.	Ye, L., Sun, P.-H., Malik, M. F. A., Mason, M. D. & Jiang, W. G. Capillary morphogenesis gene 2 inhibits growth of breast cancer cells and is inversely correlated with the disease progression and prognosis. *J. Cancer Res. Clin. Oncol.* **140**, 957–967 (2014).

3.	Rouleau, C. *et al.* The systemic administration of lethal toxin achieves a growth delay of human melanoma and neuroblastoma xenografts: Assessment of receptor contribution. *Int. J. Oncol.* **32**, 739–748 (2008).

4.	Gebhardt, C., Németh, J., Angel, P. & Hess, J. S100A8 and S100A9 in inflammation and cancer. *Biochem. Pharmacol.* **72**, 1622–1631 (2006).

5.	Watson, J., Salisbury, C., Banks, J., Whiting, P. & Hamilton, W. Predictive value of inflammatory markers for cancer diagnosis in primary care: a prospective cohort study using electronic health records. *Br. J. Cancer* **120**, 1045–1051 (2019).

6.	Li, N. *et al.* Association between C4, C4A, and C4B copy number variations and susceptibility to autoimmune diseases: A meta-analysis. *Sci. Rep.* **7**, 1–9 (2017).

7.	Blume, J. E. *et al.* Rapid, deep and precise profiling of the plasma proteome with multi-nanoparticle protein corona. *Nat. Commun.* **11**, 1–14 (2020).
